# Longitudinal Observations of the Effects of Ischemic Stroke on Binaural Perception

**DOI:** 10.1101/2023.10.12.561878

**Authors:** Anna Dietze, Peter Sörös, Henri Pöntynen, Karsten Witt, Mathias Dietz

## Abstract

Acute ischemic stroke, characterized by a localized reduction in blood flow to specific areas of the brain, has been shown to affect binaural auditory perception. In a previous study conducted during the acute phase of ischemic stroke, two tasks of binaural hearing were performed: binaural tone-in-noise detection, and lateralization of stimuli with interaural time- or level differences (Dietze et al., 2022, Front. Neurosci. 16:1022354). Various lesion-specific, as well as individual, differences in binaural performance between patients in the acute phase of stroke and a control group were demonstrated. For the current study, we re-invited the same group of patients, whereupon a subgroup repeated the experiments during the subacute and chronic phases of stroke. Similar to the initial study, this subgroup consisted of patients with lesions in different locations, including cortical and subcortical areas. At the group level, the results from the tone-in-noise detection experiment remained consistent across the three measurement phases, as did the number of deviations from normal performance in the lateralization task. However, the performance in the lateralization task exhibited variations over time among individual patients. Some patients demonstrated improvements in their lateralization abilities, indicating recovery, whereas others’ lateralization performance deteriorated during the later stages of stroke. Notably, our analyses did not reveal consistent patterns for patients with similar lesion locations. These findings suggest that recovery processes are more individual than the acute effects of stroke on binaural perception. Individual impairments in binaural hearing abilities after the acute phase of ischemic stroke have been demonstrated and should therefore also be targeted in rehabilitation programs.

## 1 Introduction

In ischemic stroke, the blood flow through the brain is suddenly disrupted by an acute blockage of blood vessels. The brain regions that are supplied with oxygen and nutrients by these blocked blood vessels can be damaged, which can result in various symptoms, including motor deficits, cognitive decline, and sensory impairments. Consequently, stroke is one of the leading causes of disability in Germany (Robert-Koch Institute, 2017) and globally the second leading cause of death (World Health Organization, 2020). After stroke, recovery of overall functional ability, including the sensory domain, is typically observed, with the majority of recovery occurring within the first weeks to months, with only reduced recovery thereafter (e.g., Skilbeck et al., 1983; Lee et al., 2015). In addition to compensatory behaviors, it has been shown in various studies that adaptive and maladaptive neuroplasticity allows the nervous system to respond to intrinsic and extrinsic stimuli by a reorganization of its structure, function, and connections (Cramer et al., 2011).

Simple hearing tasks, such as pure-tone detection, are rarely affected by stroke. According to Häusler and Levine (2000), this is due to two factors: First, many structures early in the auditory pathway, such as the cochlear nucleus, the inferior colliculus, and the medial geniculate body, have multiple sources of blood supply. Second, the bilateral structure and hemispheric crossings at and above the level of the superior olivary complex create redundancy of information in the two brain hemispheres. This leads to the phenomenon that abnormalities in hearing tasks (with the exception of speech understanding and production, which is predominantly represented in the left hemisphere) often only occur in bilateral lesions (Häusler and Levine, 2000).

Contralesional impairments, such as difficulties in moving the right arm after a lesion of the left motor cortex, are well-known symptoms of stroke (and are used to identify stroke without brain imaging). Also, visuospatial impairments, especially the phenomenon of hemispatial neglect, are often observed after right-hemispheric ischemic stroke (Heilman and Valenstein, 1979). Rarely, studies of the phenomenon of neglect also included investigations of its effects on the perception of sounds (Guilbert et al., 2016).

Binaural hearing allows listeners to accurately localize sound sources and to benefit from spatially separated target and distractor sounds by using interaural differences in time (ITD) and interaural level differences (ILD). This requires an integration of the signals captured at the left and right ear, which occurs for the first time in the brainstem, more precisely in the superior olivary complex, with representations at higher stages (e.g., Goldberg and Brown, 1969). Accordingly, lesions in different areas of the brain can interrupt binaural processing, as shown in several preceding studies (e.g., Jenkins and Masterton, 1982; Bisiach et al., 1984; Aharonson et al., 1998; Spierer et al., 2009). In general, less is known about stroke-induced effects on binaural hearing abilities compared to other modalities, especially to the visual domain. The aforementioned studies demonstrated difficulties in spatial hearing tasks, each for a specific lesion location. Including patients with different lesion locations in the same study revealed very diverse effects on binaural hearing for lesions at different locations (Dietze et al., 2022).

In the free field, ILD, ITD, and spectral cues lead to spatial perception (localization) of sound sources, because they depend on the incidence angle and the frequency of the sound (e.g., Thompson, 1882). In headphone experiments, it is possible to manipulate ILD and ITD cues independently. A presentation of unnatural ITD - ILD combinations or ILDs without the natural frequency dependence leads to an intracranial perception of sound sources that can be either in the center of the head or perceived closer to one of the ears, which is referred to as lateralization. These unnatural modifications of auditory inputs can be used to investigate ITD- or ILD-specific processing deficits that might not be detectable when both cues are congruent and spectral cues are present, as in free-field sound-localization experiments.

In psychoacoustic experiments with healthy participants, altered binaural cues lead to distorted sound source localization at first, but they partially adapt within a few days of exposure (see Wright and Zhang, 2006 for a review). One example is the 50% reduction of localization bias caused by an artificially introduced ITD bias (Javer and Schwarz, 1995). A larger ITD bias in patients with a hearing aid in one ear and a cochlear implant in the other ear, however, cannot be compensated for by the auditory system and requires a technical latency compensation (Angermeier et al., 2023). Another method is the unilateral wearing of earplugs that leads to localization distortions toward the open ear at first, but decreases over the course of five days of extensive training (e.g., Florentine, 1976; Butler, 1987). Also, experiments conducted under water showed training effects on sound source localization with altered binaural cues (e.g., Feinstein, 1973). Due to the higher speed of sound and a reduced head-shadow effect, ITDs and ILDs are diminished under water. Importantly, only specific acoustic features appear to be relearned. Re-learning does not always generalize to non-trained stimuli (reviewed by Keuroghlian and Knudsen, 2007) and adaptations to altered spatial cues are faster with training, compared to exposure (Mendonça, 2014). Following the various findings of at least partial adaptation to altered binaural information in healthy participants, and the reports of functional recovery following stroke (e.g., Skilbeck et al., 1983; Cramer et al., 2011; Lee et al., 2015), partial or full recovery of binaural perception is also expected for clinical populations such as the patients with mild symptoms of stroke, as described in Dietze et al. (2022).

To the best of our knowledge, there has been no study of the longitudinal development of binaural hearing performance after ischemic stroke. Most studies were conducted in the chronic phase of stroke, and only a few in the acute phase, but none included more than one measurement. Exploring the longitudinal effects of ischemic stroke on binaural hearing is important for two reasons: First, to gain a deeper basic knowledge of binaural processing, and second, to understand the mechanisms of recovery, and thus in consequence being able to improve auditory rehabilitation after stroke.

In this study, we aimed to quantify the effects of ischemic stroke on binaural perception in a population of stroke patients with only mild symptoms and with different lesion locations via longitudinal measurements in the acute, subacute, and chronic phase of stroke. Recovery of binaural performance was hypothesized for this group of patients whose lateralization was impaired in the acute phase of stroke, despite almost no clinical signs of stroke.

## 2 Methods

The experimental methods used in this study are identical to those reported in Dietze et al. (2022). Whenever applicable, only a summary is given here. Where all details are necessary, we quote directly from Dietze et al. (2022).

### 2.1 Measurement Phases

The study consisted of three measurements. The same experiments were conducted on all three appointments. The first measurement was in the acute phase of stroke, on average 5 days after stroke onset (results were presented in Dietze et al., 2022). The second was in the subacute phase of stroke, on average 30 days after stroke onset. The third measurement was in the chronic phase of stroke, on average 306 days after stroke onset.

### 2.2 Participants

The pool of participants is identical to those measured in the study by Dietze et al. (2022): “In total, 50 stroke patients (mean age of 63 years, SD: 14 years, 20 female, 30 male) and 12 control subjects (mean age of 61 years, SD: 14 years, 9 female, 3 male) participated after passing audiometric and cognitive assessments (see sections 2.5.1 and 0 for details) and providing written informed consent. Participants that had a stroke will be referred to as patients, whereas those participating in the control group will be referred to as control subjects. The study was approved by the Medical Research Ethics Board of the University of Oldenburg, Germany. The stroke patients were recruited in the stroke unit of the Evangelisches Krankenhaus, Oldenburg, Germany. Only those patients participated, who could understand and produce speech, who were mobile and in a general stable condition, and able to complete the different tasks despite their recent stroke. Exclusion criteria were additional neurological diseases or a pure-tone average of 40 dB HL or more (see section 2.5.1). The control group was age-matched and followed the same exclusion criteria.” A subset of 15 patients did the experiments in all three measurement phases (acute, subacute, and chronic).

#### 2.2.1 Acute Phase Measurements

Of the 50 patients who participated in the acute phase measurements as reported in Dietze et al. (2022), only 31 also participated in at least one of the later measurements. All further analyses for the acute phase measurements are based on these 31 patients. They were tested in a quiet room at the Evangelisches Krankenhaus, Oldenburg, Germany, on average 5 days (range: 2 – 8 days) from stroke onset. The National Institute of Health Stroke Scale revealed that the patients only suffered from minor stroke symptoms. The scores of the patients ranged from 0 to 5 points (the maximum possible score is 42, with higher scores indicating worse signs and symptoms of ischemic stroke).

#### 2.2.2 Subacute Phase Measurements

A subset of 24 patients (mean age of 58 years, SD: 15 years, 9 female, 14 male) participated in the experiments in the subacute phase of stroke. They were tested in a quiet room at the Rehazentrum Oldenburg, Germany, on average 30 days (range: 23-39 days, 13 days for S27) after stroke onset.

#### 2.2.3 Chronic Phase Measurements

A subset of 22 patients (mean age of 65 years, SD: 14 years, 6 female, 16 male) participated in the experiments in the chronic phase of stroke. They were tested in an acoustically shielded chamber at the University of Oldenburg, Oldenburg, Germany, on average 306 days (range: 216-391 days, 500 days for S51) after stroke onset.

### 2.3 General assessment

Identical to the study in the acute phase (Dietze et al., 2022), the Montreal Cognitive Assessment (MoCA, Nasreddine et al., 2005) and the short version of Beck’s Depression Inventory (BDI, Beck et al., 2013) were conducted in the subacute and chronic phase, to screen for mild cognitive impairment or dementia and to quantify the severity of possible depression.

### 2.4 Magnetic resonance imaging

Lesion location and lesion volume were extracted from magnetic resonance imaging (MRI), as explained in Dietze et al. (2022). In summary, after brain-extraction, linear registration to the structural template, provided by the Montreal Neurological Institute (MNI 152) was performed for the fluid-attenuated inversion recovery images and the lesion masks. Finally, we checked for possible overlap of the MNI-registered stroke lesions with brain areas belonging to the auditory pathway.

### 2.5 Psychoacoustic experiments

Custom Matlab scripts using the psychophysical measurement package AFC (Ewert, 2013), were used to generate and reproduce the stimuli. Closed headphones with passive sound attenuation (HDA300, Sennheiser electronic GmbH, Wedemark, Germany), driven by an external sound card (UR22mkII, Steinberg Media Technologies GmbH, Hamburg, Germany) were used for all psychoacoustic experiments, with the exception of audiometric testing in the chronic phase. For these measurements, the system specified in section 2.5.1 was used.

#### 2.5.1 Audiometry

Identical to the acute phase measurements described in Dietze et al. (2022), pure-tone audiometric thresholds were also measured in the subacute phase for a restricted set of frequencies (500 Hz, 1000 Hz, 3000 Hz). The same hardware as for the other experiments (see section 2.4) was used. In the chronic phase measurement, a clinical pure-tone audiometric air-conduction test ranging from 125 Hz to 8000 Hz was performed (Equinox 2.0, Interacoustics, Middelfart, Denmark).

The pure-tone average over the three frequencies was calculated individually for the left ear (PTA3 L) and right ear (PTA3 R) and averaged (PTA3). To estimate the asymmetry of pure tone hearing loss, the difference between left and right PTA3 (PTA3 asymmetry) was calculated.

#### 2.5.2 Tone-in-noise detection

The tone-in-noise detection experiment was conducted as described in Dietze et al. (2022): “The participants were presented with three intervals containing 500-ms bursts of octave-wide white noise centered around 500 Hz (333 Hz – 666 Hz). The stimuli were gated with 20-ms raised cosine onset and offset ramps. The intervals were separated by 300-ms silent gaps. In one of the three intervals, an additional 500-Hz pure-tone of 420 ms duration was added and temporally centered in the noise. The tone had the same ramp parameters as the noise, but its onset was 40 ms later than the noise.

Similarly, the tone offset was 40 ms before the noise offset. The participants’ task was to detect the deviating interval (the one containing the tone) and to press key number ‘1’, ‘2’, or ‘3’ on a computer keyboard, indicating whether the first, second, or third interval was the odd one. The tone was either interaurally in phase with the noise (condition N_0_S_0_) or had an interaural phase difference of π (condition N_0_S_π_). The experiment started without any training and with two runs of the N_0_S_π_ condition. This was followed by one run of the N_0_S_0_ condition. The noise was presented with 60 dB sound-pressure level (SPL). The level of the tone was initially 65 dB SPL in the N_0_S_0_ condition and 50 dB SPL in the N_0_S_π_ condition. The level varied according to a one-up, three-down procedure, with a step size of 4 dB up to the second reversal, and a step size of 2 dB for the remaining 8 reversals, converging to 79.4% correct thresholds. Thresholds are calculated as the average of the last 8 reversals. If the staircase track hit the maximum tone level of 80 dB SPL during a measurement, re-instructions on how to perform the task were provided. If this did not lead to improvements in task performance, the run was stopped and marked as invalid. No feedback was given during the runs. The binaural masking level difference (BMLD) was calculated from the threshold difference between N_0_S_0_ and the better of the two N_0_S_π_ runs.”

#### 2.5.3 Lateralization

The lateralization task and the calculation of variables for quantitative description of the lateralization pattern were also carried out as reported in Dietze et al. (2022): “For the lateralization task, again, a one-octave wide white noise, centered around 500 Hz with an interaural difference in either level or time was presented. The stimuli were generated by copying the same noise sample to both channels and then applying the interaural difference in time or level. The task was to indicate where the sound was perceived inside the head. Responses were given by pressing one of the horizontally aligned numbers ‘1’ to ‘9’ on a computer keyboard, above the letter keys. The participants were instructed to press ‘1’ when the sound was heard on the very left side of their head, ‘5’ for sounds perceived in the center of the head and ‘9’ for the very right side. For possible intracranial positions between the center and the two extremes, the participants were asked to press the respective number ‘2’,’3’,’4’,’6’,’7’ or ‘8’ on the keyboard. For visual guidance, a template with a schematic drawing of a head indicated the positions of the ears and the center relative to the response buttons. The template covered all of the keyboard except for the numbers ‘1’ to ‘9’. The duration of the stimuli was 1 s, gated with cosine ramps of 10 ms duration and presented at 70 dB SPL. ITDs ranging from −600 µs to 600 µs in steps of 200 µs, and two ITDs outside the physiological range (−1500 µs and 1500 µs), were presented. The ILDs ranged from −12 dB to 12 dB in steps of 4 dB. The level of the left- and right-ear signals was changed without changing the overall energy by applying the formula presented in Dietz et al. (2013). In addition, monaural stimulation of the left ear and right ear was tested. Each stimulus was presented six times in random order. The diotic stimulus (zero ITD/ILD) was presented eight times. To ensure one common reference system for both types of interaural differences, ILD and ITD stimuli were presented interleaved. In contrast to the investigations by Furst et al. (2000), no training and no center reference were provided in our study. The response to the first trial of each stimulus was not used in further analyses.

Several variables for quantitative description of the lateralization pattern were calculated:

A linear fit to the three left-favoring and right-favoring stimuli, individually for ILD stimuli (−12 dB, −8 dB, −4 dB and 4 dB, 8 dB, 12 dB) and ITD stimuli (−600 µs, −400 µs, −200 µs and 200 µs, 400 µs, 600 µs) was used to describe the steepness of the participants’ lateralization percept (*ILD L slope, ILD R slope, ITD L slope, ITD R slope*). The logarithmic ratio of the left and right slope (*ILD slope ratio, ITD slope ratio, e.g., ILD slope ratio = log(ITD slope L / ILD slope R))* indicates an asymmetric steepness of the two sides.

Variables that inform about side biases in the responses were calculated: The mean of the responses to all ITD or all ILD stimuli (*ITD mean, ILD mean*) and the mean of the fit to left-favoring and right-favoring stimuli (*ITD L fit, ITD R fit, ILD L fit, ILD R fit*) were calculated. Furthermore, the mean of those stimuli that were perceived as being in the center of the head (when key ‘5’ was pressed), was calculated for ILD and for ITD stimuli (*ITD center, ILD center*). The so-called *diotic percept* was the mean of the responses given for the zero ILD/ITD stimuli.

Another feature of the lateralization data is its variability. For this, the standard deviation for zero ILD/ITD was calculated (*diotic std*), as well as the mean of the standard deviations of the responses to each ILD stimulus (excluding the monaural stimulation, *ILD std*), each ITD stimulus (*ITD std*) and the mean standard deviation of the left-favoring and right-favoring stimuli independently (*ITD L std, ITD R std, ILD L std, ILD R std*). Their logarithmic ratios (*ITD std ratio, ILD std ratio)* can indicate differences in the variability of left-favoring and right-favoring stimuli.

The maximal range of lateralization was calculated by the difference of the maximally lateralized responses given for ITDs within the physiological range (*ITD range*), and for all ILDs excluding monaural stimulation (*ILD range*). The logarithmic ratio of the ranges obtained with ILD and ITD stimuli (*range ratio*) informs about differences in the ranges perceived using the two types of stimuli.

The perception of the monaural left and right (*mon left, mon right*), and the ITDs of ± 1500 µs (*neg 1500, pos 1500*) was only evaluated in terms of the mean response to these stimuli.

For all the variables, values within the interval of 1.5 times the standard deviation around the control group mean were considered to be normal. As we did not want to overemphasize possible asymmetries of the left and right side of individual control subjects, we also added the mirrored control data before calculating the mean and standard deviation. With this, the mean was not biased by individual asymmetries and the standard deviation remained unaffected. We verified that adding the mirrored data did not change the results substantially from those obtained without adding the mirrored responses to the data set. Whenever values of the calculated variables are reported, they are in the unit of response keys (a difference of one response button corresponds to 1/8 of the distance between the two ears), except for the variables describing the goodness of fit and the ratios.”

On average, four divergences from the control can be expected even in a normally performing participant, because 13% of Gaussian distributed data are outside the range of −1.5 to +1.5 standard deviations, which corresponds to 4 out of 31 possible divergences. Participants with no more than four divergences from the control group are therefore defined as having normal lateralization.

To quantify the change in lateralization performance, the change in each of the lateralization metrics from one phase to the other is expressed in units of standard deviation of the respective metric in the control group. Positive changes are those where the metric values in the later measurements approach the mean of the control group, and negative changes are those where the metric values in the later measurements diverge more from the control group’s mean.

## 3 Results

### 3.1 General assessment

Mean values and standard deviations of the non-auditory testing and the audiometric results of the control group and the stroke groups at the three measurement phases (acute, subacute, and chronic) are shown in Table 1. According to statistical tests (two-sample t-tests, statistics shown in the right-most column), the control group and the chronic phase group did not differ in age, not in their pure-tone average over the three tested frequencies, and also not in the absolute asymmetry of their left and right PTA3. The scores for the short form of Beck Depression Inventory (BDI) also did not differ significantly between these two groups. In the cognitive screening test (MoCA) however, significantly lower scores were obtained for the chronic phase stroke group compared to the control group.

**Table 1.**
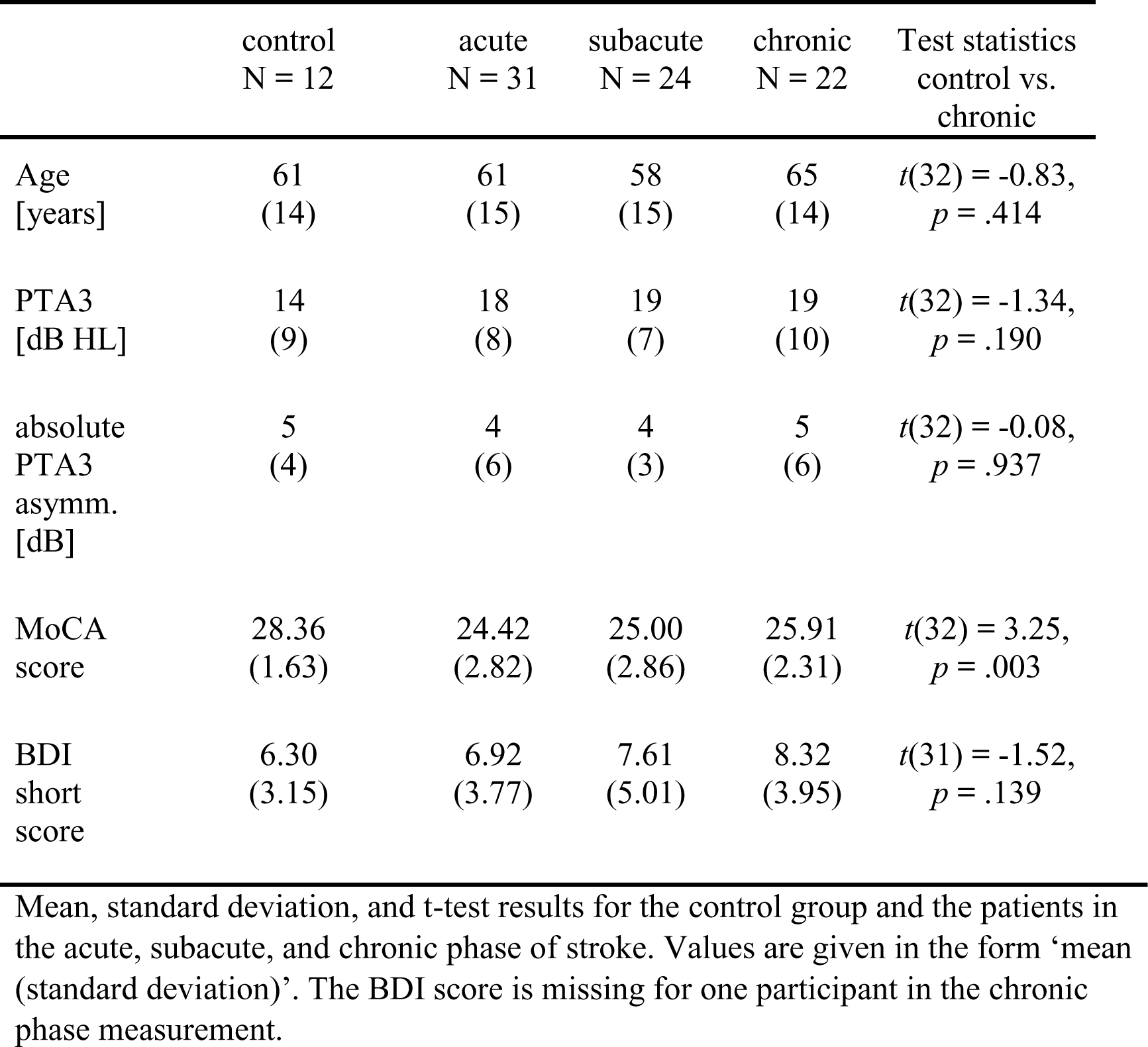
General assessment results.

|  | control<br>N = 12 | acute<br>N = 31 | subacute<br>N = 24 | chronic<br>N = 22 | Test statistics<br>control vs.<br>chronic |
| --- | --- | --- | --- | --- | --- |
| Age<br>[years] | 61<br>(14) | 61<br>(15) | 58<br>(15) | 65<br>(14) | $t(32) = -0.83$,<br>$p = .414$ |
| PTA3<br>[dB HL] | 14<br>(9) | 18<br>(8) | 19<br>(7) | 19<br>(10) | $t(32) = -1.34$,<br>$p = .190$ |
| absolute<br>PTA3<br>asymm.<br>[dB] | 5<br>(4) | 4<br>(6) | 4<br>(3) | 5<br>(6) | $t(32) = -0.08$,<br>$p = .937$ |
| MoCA<br>score | 28.36<br>(1.63) | 24.42<br>(2.82) | 25.00<br>(2.86) | 25.91<br>(2.31) | $t(32) = 3.25$,<br>$p = .003$ |
| BDI<br>short<br>score | 6.30<br>(3.15) | 6.92<br>(3.77) | 7.61<br>(5.01) | 8.32<br>(3.95) | $t(31) = -1.52$,<br>$p = .139$ |

Mean, standard deviation, and t-test results for the control group and the patients in the acute, subacute, and chronic phase of stroke. Values are given in the form ‘mean (standard deviation)’. The BDI score is missing for one participant in the chronic phase measurement.

### 3.2 Audiometry

The PTA3, calculated over the three frequencies for each of the three measurements, showed that the pure-tone hearing thresholds were comparable for the three measurement phases and the two audiometric measurement procedures. A PTA3 of 20 dB HL or higher was reached in 39%, 35%, and 38% of the patients in the acute, subacute, and chronic phase measurements, respectively. Only taking the 15 patients that participated in all three measurements into account, a repeated measures ANOVA revealed no significant effect of the three measurement phases (acute, subacute, chronic: *F*(2, 28) = 1.82, *p* = .18, *η*² = .007) or side (left, right: *F*(1, 14) = 0.32, *p* = .58, *η*² = . 001), nor an interaction of measurement phase and side on those participants’ PTA3 (*F*(2, 28) = 2.52, *p* = .09, *η*² = . 004).

In Figure 1A-C it can be seen that in all three measurement phases there is a statistically insignificant trend towards worse hearing thresholds for the first ear measured (left ear in acute and subacute phase, right ear in chronic phase; acute: *t*(30)=1.55, *p* =.13, subacute: *t*(23)=0.57, *p* = .57, chronic: *t*(21)=-0.07, *p* = .95).

**Figure 1.**
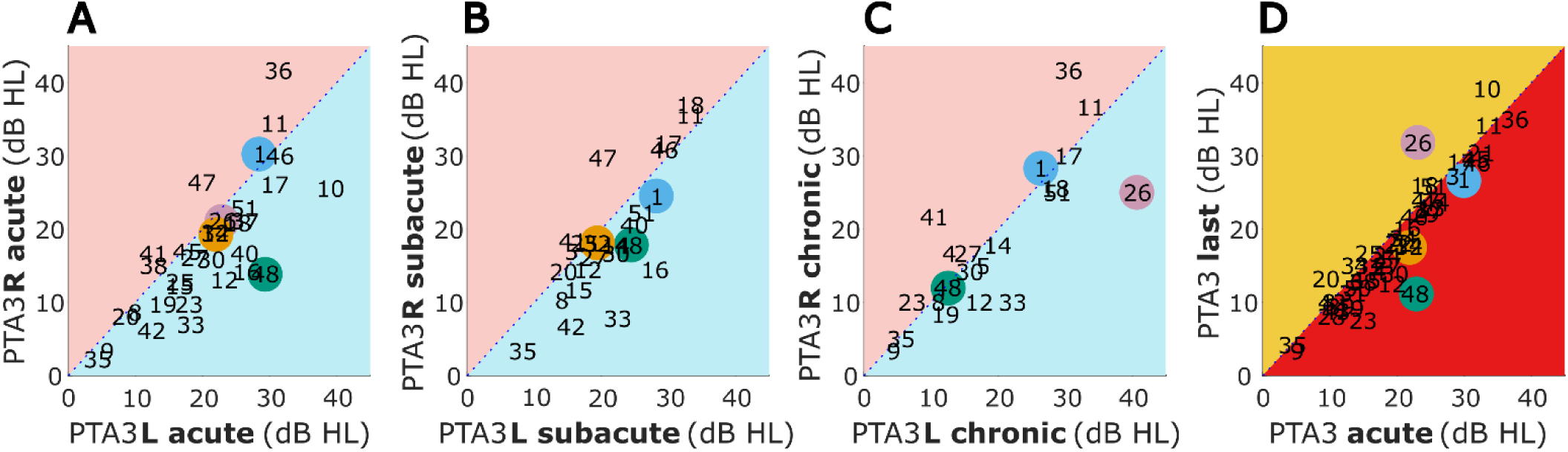
Mean pure-tone audiometric thresholds for 500, 1000, and 3000 Hz (PTA3) for the left vs. right side for the acute, subacute, and chronic phase (panels **A-C**) and for acute vs. last measurement (panel **D**). Selected patients are highlighted by the color coding used throughout the figures.

A comparison of the PTA3 of the acute phase measurement and the last measurement is presented in Figure 1D. Only for a few patients (S10, S20, S23, S26, and S48) did the PTA3 of the two measurements differ by more than 5 dB. Of these five individuals, two had improved and three had deteriorated hearing thresholds. In general, no significant difference in hearing thresholds of the acute and the last measurement were observed (t(30) = −0.70, *p* = 0.49).

### 3.3 Correlation analyses

Partial correlations between the variables *days after stroke*, *PTA3*, *age*, and the results of *MoCA* and *BDI*, while controlling for the remaining three variables, are shown in Table 2. They revealed a weak positive correlation of *days after stroke* and *MOCA*. *Age* and *PTA3* had a moderate positive correlation and *age* and *MOCA* had a weak negative correlation. None of the other variables were significantly correlated, when controlling for the remaining variables. A Spearman rank correlation was conducted, because not all variables, especially not the variable *days after stroke*, were normally distributed.

**Table 2.**
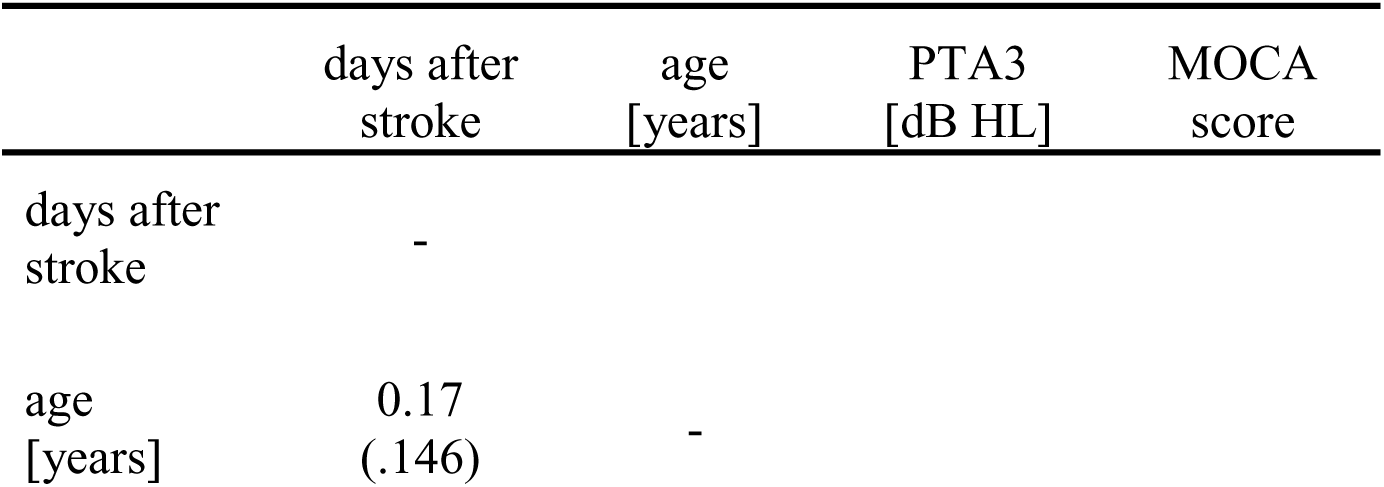
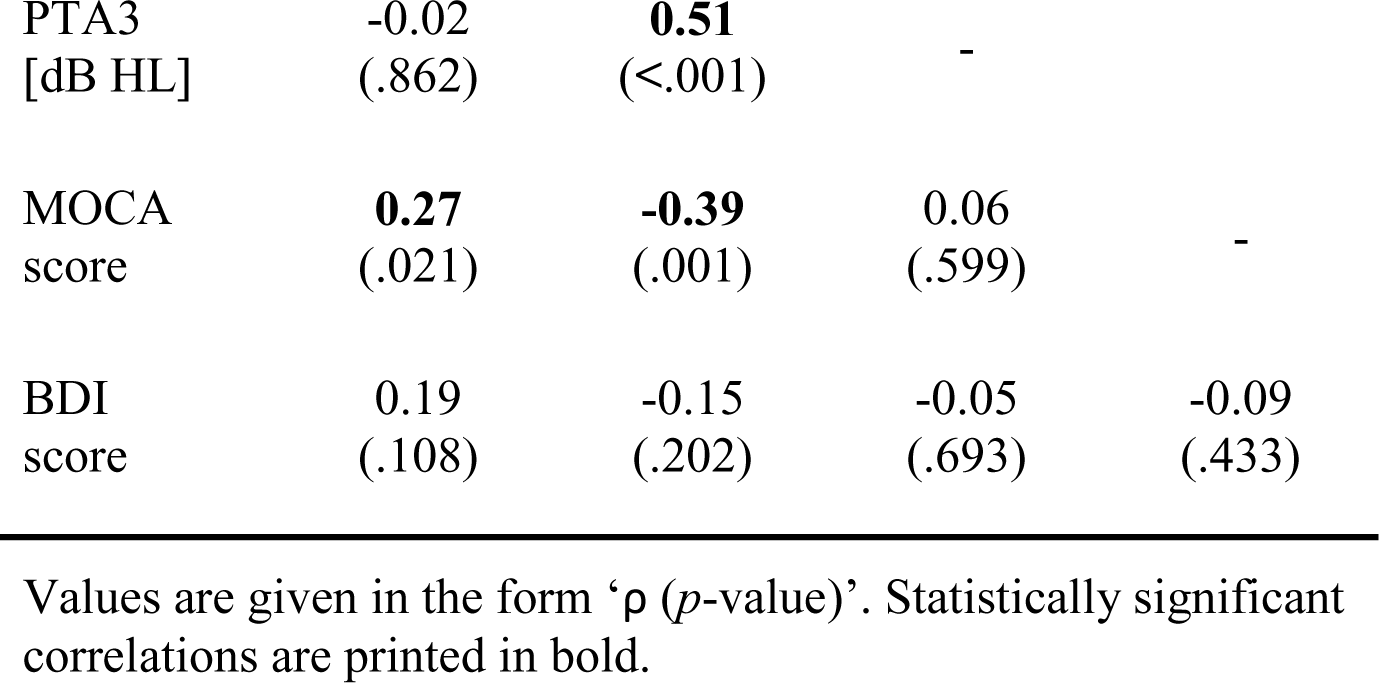
Partial Spearman Correlation.

### 3.4 Tone-in-noise detection

In the acute phase measurements, 29 of 31 stroke patients produced converging tracks in both conditions of the tone-in-noise detection task (N_0_S_π_ and N_0_S_0_), allowing the BMLD from the difference between the N_0_S_π_ and N_0_S_0_ thresholds to be calculated. The same was true for 11 of the 12 control subjects, 23 of 24 patients in the subacute and all 22 patients in the chronic phase. Of those patients with non-converging tracks in the acute phase measurement, S20 participated again in the subacute phase and S18 in the subacute and chronic phase. In the later measurement, S20 had a BMLD of 12 dB. S18 did not produce converging tracks in the subacute phase, but had a BMLD of 15 dB in the chronic phase measurement.

The normal values of BMLD, as defined by the mean ± 1.5 times the standard deviation of the control group results, ranged from 7.5 dB to 20.1 dB. Of those participants that produced convergent tracks, a BMLD of 7.5 dB or more was measured in 27 of 29 patients in the acute phase, in all 24 patients in the subacute phase, and 21 of 22 patients in the chronic phase (see Figure 2).

**Figure 2.**
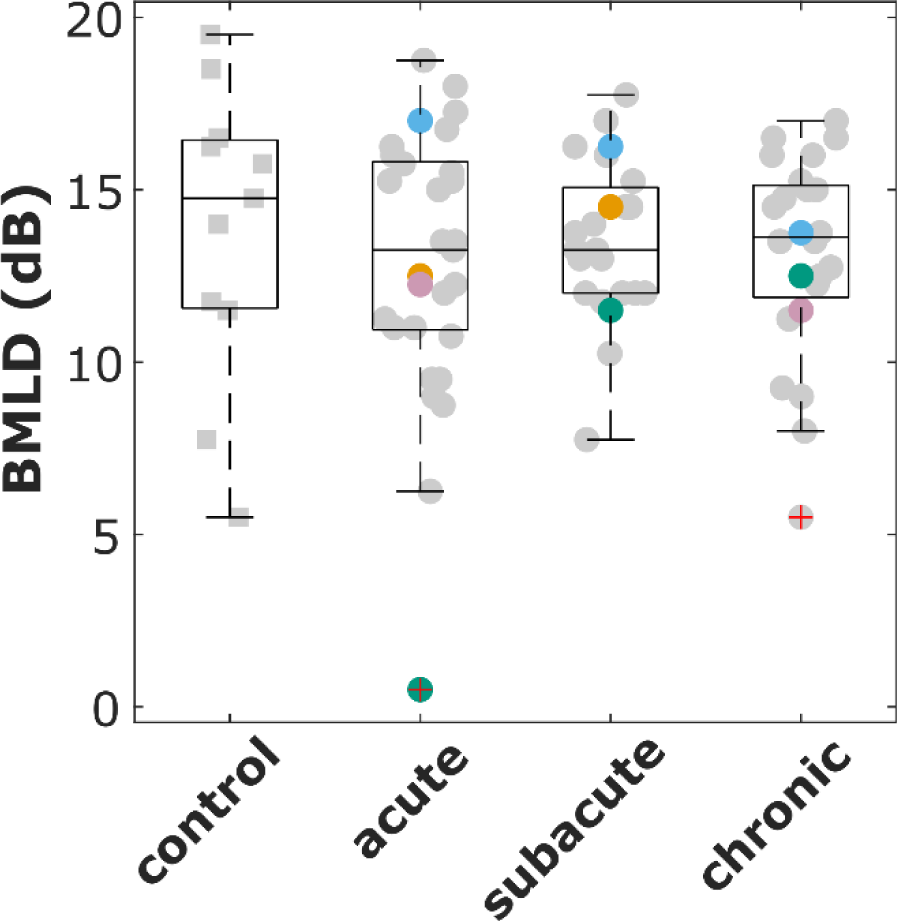
Binaural masking level difference (BMLD) for the control group and all patients that participated in the acute, subacute, and chronic phase measurements. Selected patients are highlighted by the color coding used throughout the figures.

A repeated-measures ANCOVA with the covariates MOCA and BDI was conducted for the 15 patients that both produced converging tracks and participated in all three measurements. It revealed no significant effect of the three measurement phases (acute, subacute, chronic) on the participants’ BMLD. Mauchly’s test indicated that the assumption of sphericity had been violated, therefore Greenhouse–Geisser corrected tests are reported: *F*(1.33, 17.25) = 0.18, *p* = .75, *η*² = . 005.

### 3.5 Lateralization

All control subjects and all patients completed the lateralization task in each of the measurements in which they participated. Normal lateralization (with no more than four divergences from the control group) was found for 32% of the patients in the acute, 71% in the subacute, and 41% in the chronic phase of stroke. The lateralization patterns of four selected patients (details are given in Figure 3A and Table 3) are shown in Figure 3B-E. These patients were selected, because they represent well the variety of lateralization patterns and their changes from the acute phase to the later phases. The lateralization results of all other participants can be found in the supplementary data.

**Figure 3.**
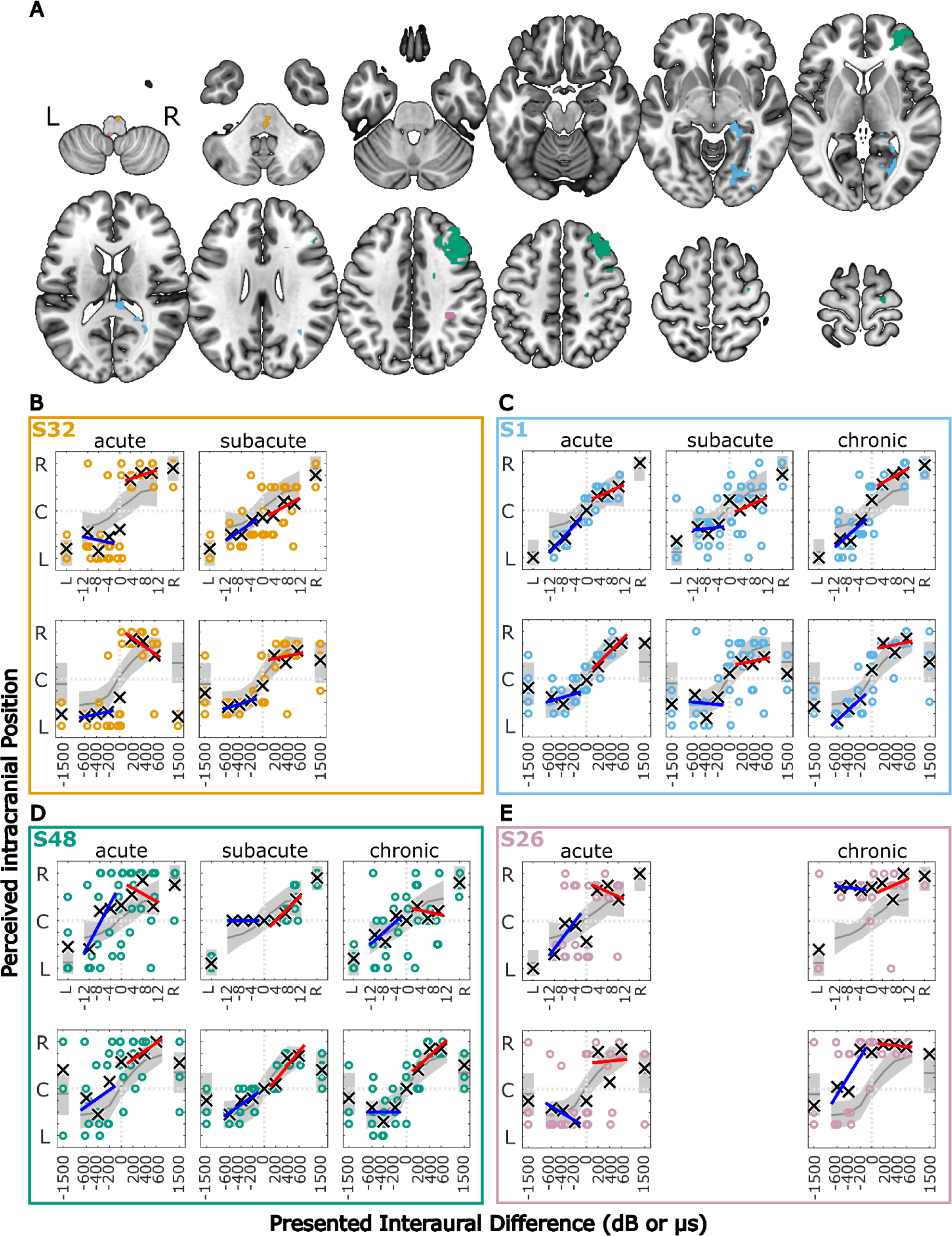
Lesion locations overlaid on axial slices of the MNI152 template (**A**). Lesion group, lesion volume and additional information is given in Table 3. **B-E**: Results of the lateralization task for four selected stroke patients in the three measurement phases. The colored symbols represent the responses given to the individual trials of the same stimulus, except for the discarded first trial. The black crosses indicate the means of the given responses. The red and blue lines represent linear fits to right-favoring and left-favoring stimuli, respectively. The gray line and shaded area indicate the mean and the 1.5 times standard deviation interval around the mean response of the control subjects. Selected patients are highlighted by the color coding used throughout the figures.

**Table 3.**
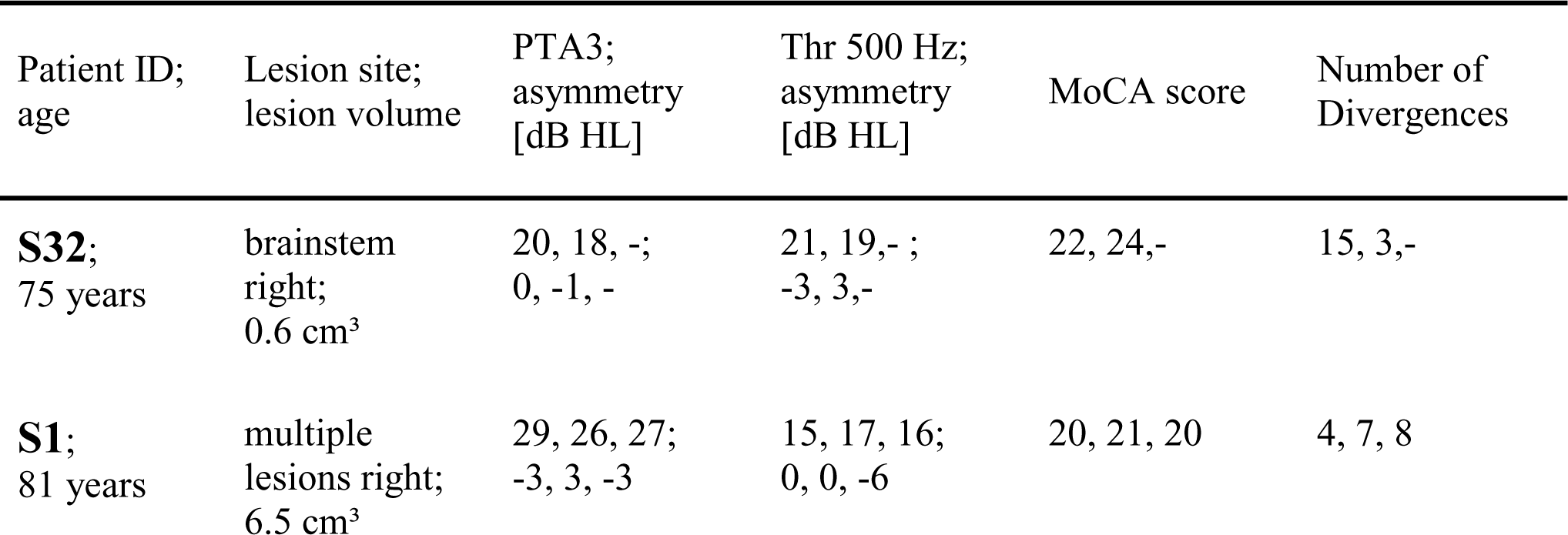
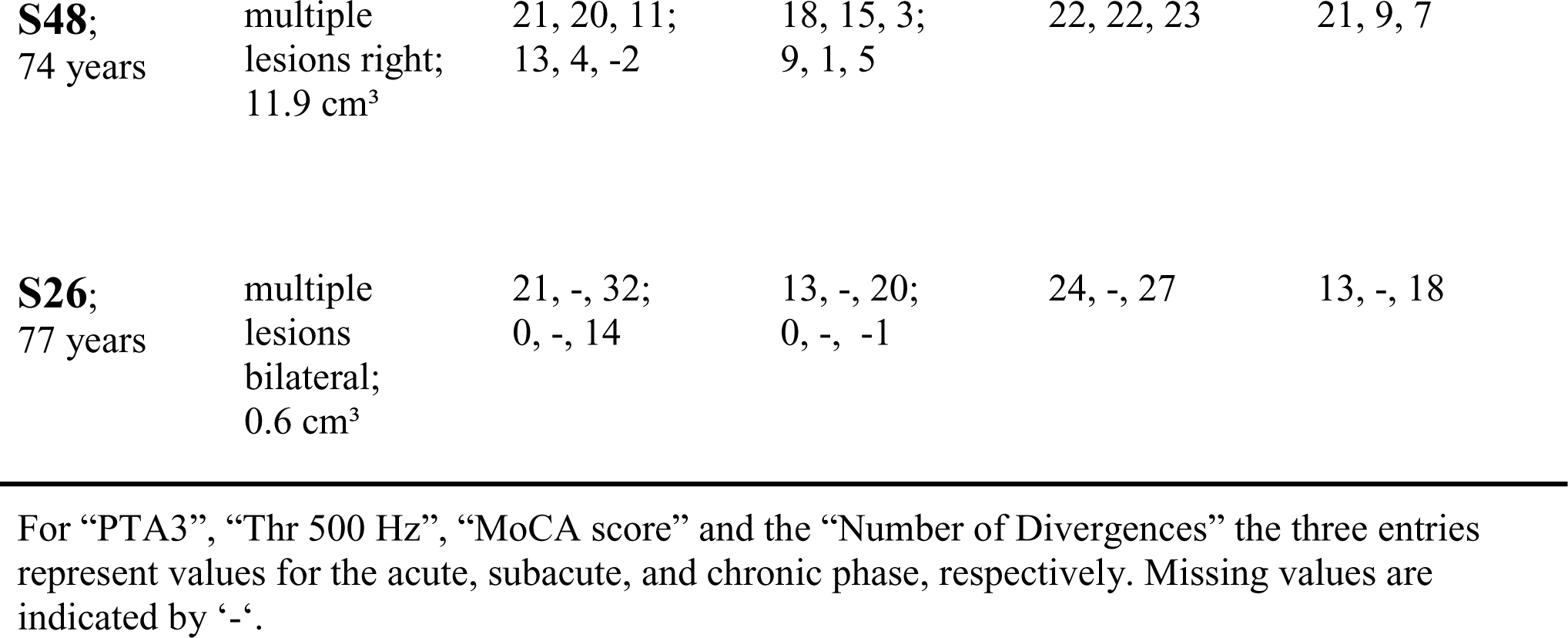
Additional information for the four selected stroke patients.

| Patient ID;<br>age | Lesion site;<br>lesion volume | PTA3;<br>asymmetry<br>[dB HL] | Thr 500 Hz;<br>asymmetry<br>[dB HL] | MoCA score | Number of<br>Divergences |
| --- | --- | --- | --- | --- | --- |
| **S32;**<br>75 years | brainstem<br>right;<br>$0.6 \text{ cm}^3$ | 20, 18, -;<br>0, -1, - | 21, 19, - ;<br>-3, 3, - | 22, 24, - | 15, 3, - |
| **S1;**<br>81 years | multiple<br>lesions right;<br>$6.5 \text{ cm}^3$ | 29, 26, 27;<br>-3, 3, -3 | 15, 17, 16;<br>0, 0, -6 | 20, 21, 20 | 4, 7, 8 |
| **S48;**<br>74 years | multiple<br>lesions right;<br>11.9 cm<sup>3</sup> | 21, 20, 11;<br>13, 4, -2 | 18, 15, 3;<br>9, 1, 5 | 22, 22, 23 | 21, 9, 7 |
| **S26;**<br>77 years | multiple<br>lesions<br>bilateral;<br>0.6 cm<sup>3</sup> | 21, -, 32;<br>0, -, 14 | 13, -, 20;<br>0, -, -1 | 24, -, 27 | 13, -, 18 |

For “PTA3”, “Thr 500 Hz”, “MoCA score” and the “Number of Divergences” the three entries represent values for the acute, subacute, and chronic phase, respectively. Missing values are indicated by ‘-’.

Across the three measurement phases, the metric ‘diotic percept’ that is associated with a spatial percept away from the auditory midline for stimuli without any ILD or ITD is positively correlated with the asymmetry of PTA3 (ρ = 0.446, p < 0.001).

#### 3.5.1 Single-patient observations

Patient S1 (81 years) had multiple lesions in the right hemisphere, including the occipital lobe, lingual and fusiform gyrus, hippocampus, thalamus and the corpus callosum, with a total lesion volume of 6.5 cm³ (see Figure 3A, C, and Table 3). As shown in Figure 3C, the lateralization pattern was very close to the control group in the acute phase, becoming more variable in the subacute phase. In this phase, there were left-right confusions for physically right-leading, but less for left-favoring stimuli for ILD and ITD stimuli. A slight shift towards the right side can be observed in the subacute and chronic phases. The hearing thresholds at the right ear for 500 Hz were worse by 6 dB in the chronic phase but were within normal limits for the other frequencies and other measurement phases.

In Figure 3B, side-oriented lateralization with both ILD and ITD cues in the acute phase measurement can be seen for patient S32 (75 years) who had a lesion in the right pons (total lesion volume: 0.6 cm³, covering parts of the primary auditory pathway, see Figure 3A, B, and Table 3).

The stimulus was perceived close to the ears even for small ILDs or ITDs, with a few left-right confusions for physically left-favoring stimuli. Both stimuli with unnaturally large ITDs (± 1500 µs) were perceived on the left side. In the subacute phase, the lateralization pattern resembled the control group in most aspects, including the +1500-µs stimulus now being perceived on the leading right side. This patient’s pure tone audibility stayed constant over time, with no strong asymmetry between left and right ear thresholds.

Patient S26 (77 years, see Figure 3A, E, and Table 3) had several lesions in the right hemisphere, including sulcus intraparietalis, superior temporal lobe, and anterior insula and a lesion in the dorsal left medulla oblongata (total lesion volume: 0.6 cm³). As in patient S32, this patient showed side-oriented lateralization patterns for ILD and ITD stimuli in the acute phase, but did not recover to normal lateralization in the later measurement (chronic phase). Instead, a shift of lateralization towards the right side was observed. Importantly, the pure-tone hearing thresholds of this patient were symmetric in the acute phase, but asymmetric in the chronic phase measurements, with the PTA3 of the left ear being 14 dB worse than for the right ear, but with no asymmetry at 500 Hz. Interestingly, all stimuli with ILDs (favoring either the left or the right ear) were perceived on the right side, whereas the ITDs of −400, −600, and −1500 µs were frequently perceived on the leading left side.

Marked changes in the lateralization pattern from the acute to the subacute and chronic phase can be seen in patient S48 (74 years, see Figure 3A, D, and Table 3). This patient had multiple lesions in the right hemisphere in the medial and superior frontal lobe, precentral gyrus, and several smaller right-sided white-matter lesions. The total volume of all lesion sites was 11.9 cm³. High variability for individual responses, especially to physically left-favoring stimuli, and a shift of the responses towards the right side were observed in the acute phase. In the subacute phase, the responses were much less variable, and all left-favoring ILD, but not ITD stimuli, were perceived in the center of the head. Finally, in the chronic phase, the responses were again more variable, but overall not diverging much from the control-group behavior. This patient’s PTA3 was asymmetric in the acute phase (13 dB worse in the left ear), but this was reduced to 4 and −2 dB in the subacute and chronic measurements, respectively, because of an improvement of the left ear PTA3.

#### 3.5.2 Differences to the Control Group across Measurements

The absolute number of divergences from the control group in the lateralization metrics (described in section 2.5.3) is shown for all patients in Figure 4. Values on the diagonal represent patients with the same number of divergences in two measurements (see patient S25 in panel A), whereas numbers below the diagonal represent patients with a smaller number of divergences in the later stages. More patients are below the diagonal (improved lateralization) than above the diagonal (deteriorated performance). When comparing panels A and B to panel C, it becomes clear that the number of divergences changed more from the acute to the subacute phase (on average 1.9 (SD: 5.0) fewer divergences in the later measurement) than it changed from the subacute to the chronic phase (on average 0.7 (SD: 2.9) more divergence in the later measurement). Divergences in the individual metrics for each patient and the three measurement phases can be found in the supplementary figures (Figure S1-S3).

**Figure 4.**
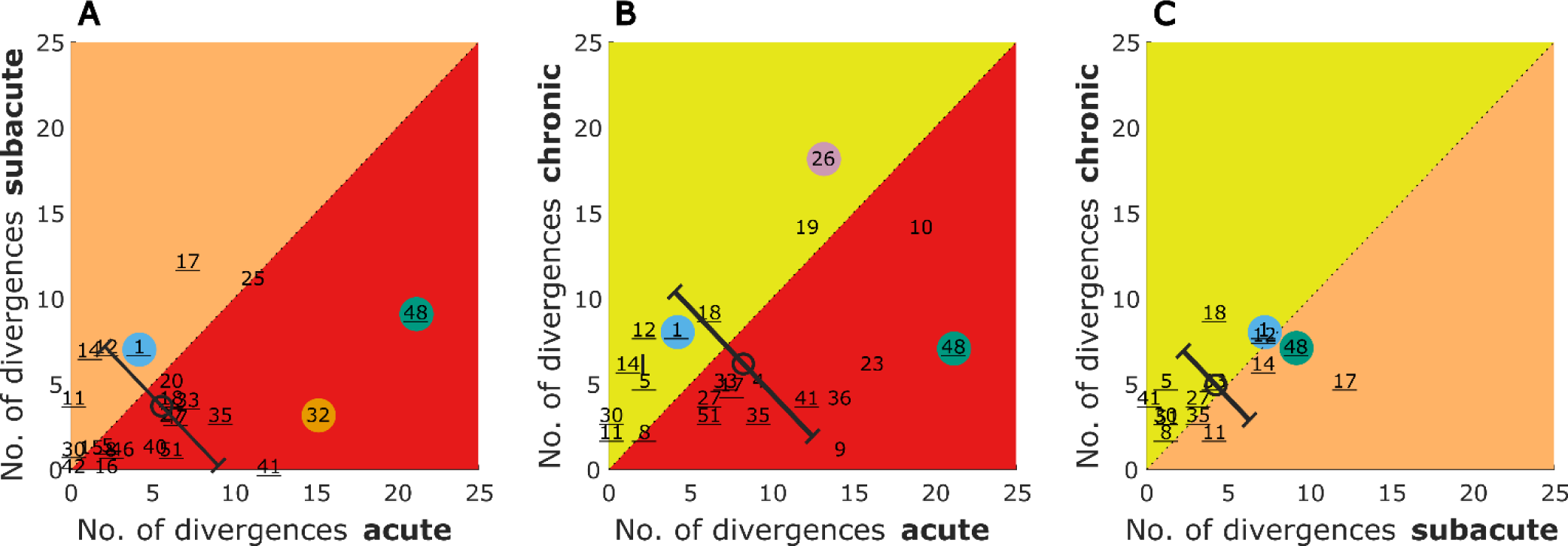
Number of divergences from the control group in the lateralization task for acute vs. subacute, acute vs. chronic, and subacute vs. chronic phase. The number refers to the patient identifier, with the number of those patients that participated in all three measurements being underlined. Selected patients are highlighted by the color coding used throughout the figures. The circles represent the mean of the patients and the error bars show the standard deviation of the change in the number of divergences from one to the other phase.

The lateralization performance of the patients S1, S26, S32, and S48 was previously described on the basis of visual inspection. These results are also reflected in the number of divergences shown for these color coded patients in Figure 4.

A repeated-measures ANCOVA with the covariates MOCA and BDI was conducted for the 15 patients that participated in all three measurements. It revealed no statistically significant effect of the three measurement phases (acute, subacute, chronic) on the number of divergences from the control group. Mauchly’s test indicated that the assumption of sphericity had been violated, therefore Greenhouse–Geisser corrected tests are reported: *F*(1.39, 19.40) = 2.38, *p* = .13, *η*² = . 086.

#### 3.5.3 Changes in Lateralization Patterns across Measurements

To be able to follow the changes in lateralization abilities across the measurement phases, we calculated the difference of the lateralization metrics in units of standard deviation of the control group across the measurements. It became clear that in the later measurement phases, some participants’ lateralization approached closer to normal behavior (i.e. the mean lateralization metric values of the control group) with respect to many metrics. For others, in the later measurements many metrics diverge even further from the control group (see in Figure 5 the blue and red squares for improved and deteriorated metric values, respectively).

**Figure 5.**
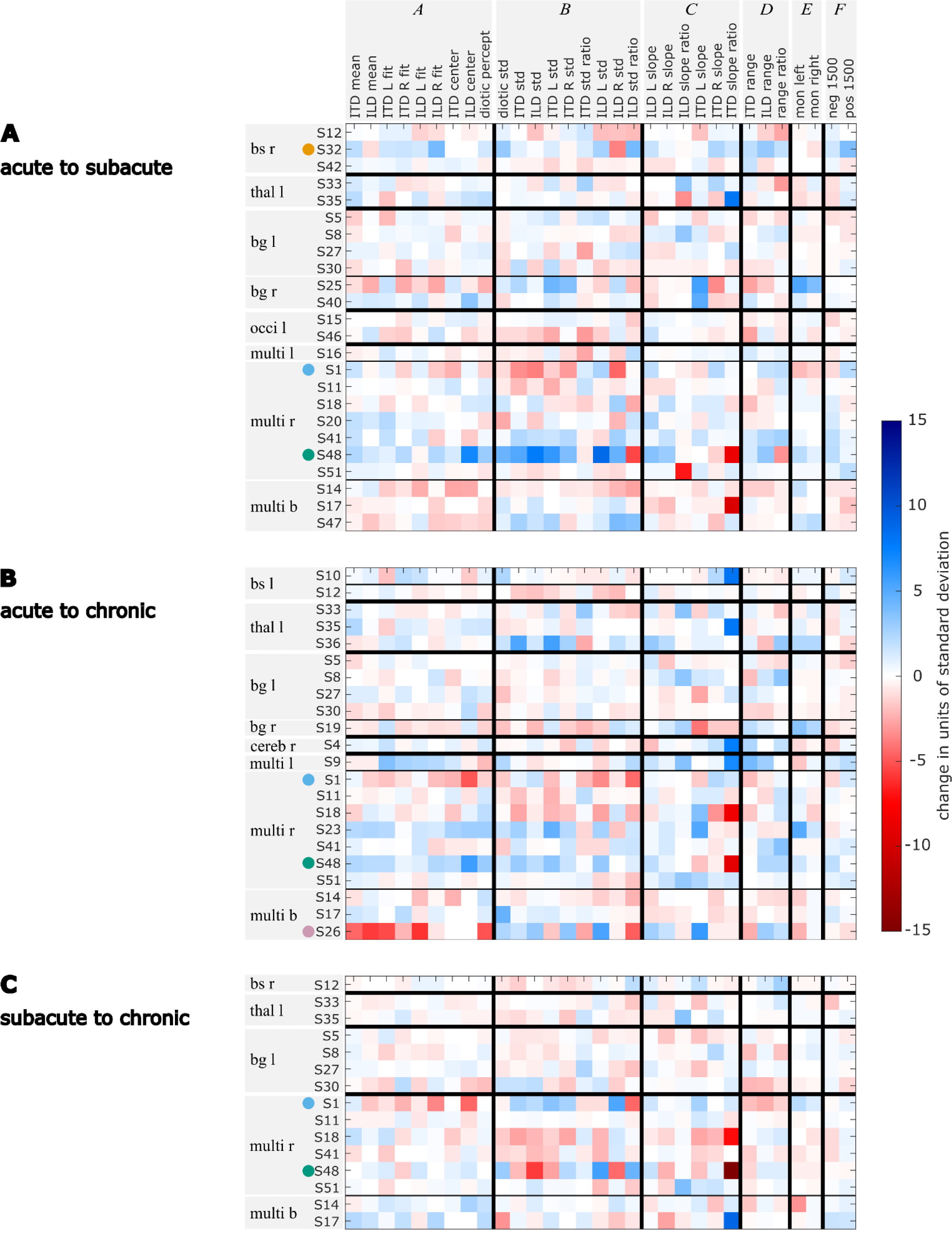
Changes in units of standard deviation of the control group for each lateralization metric for the comparisons acute vs. subacute (panel **A**), acute vs. chronic (panel **B**), and subacute vs. chronic (panel **C**). Changes that lead to the metric values being further away from the control group’s mean are marked in red, changes that lead to metric values becoming closer to the control group’s mean are marked in blue. Selected patients are highlighted by the color coding used throughout the figures.

A further examination of the example patients S1, S26, S32, and S48 revealed that the changes in their lateralization performance as described before on the basis of visual inspection of Figure 3 is reflected in the quantitative analyses presented in Figure 5: For patient S1, the temporary increase in variability in the subacute phase and the shift towards the right side that was observed in the subacute and chronic phases, is clearly reflected in the high number of deteriorated values in the respective clusters B and A in Figure 5A, B, and C. The improvements in the lateralization performance of patient S32 can be clearly observed in Figure 5A. Changes towards normal lateralization were found in cluster A, associated with shifts of the auditory space and in the metrics of the ±1500-µs stimuli (cluster F). The only marked change towards poorer metric values is present in the variability of right-sided ILD stimuli. For patient S26, a strong shift towards the right side is reflected in Figure 5B in the change of the metrics in cluster A (associated with shifts). Since physically left-leading ITD stimuli were not always shifted to the right side, the slope of these stimuli recovered relative to the acute phase, as seen in the positive change of ITD L slope. For patient S48, the observation of improved performance, especially with regard to variability and shift in the later stages, is reflected in Figure 5A and B. In panel B, the metric ‘ILD std ratio’ and the mixed effects in panel Figure 5C reveal the strong difference in the perception of left-and right-favoring ILD stimuli in the subacute phase.

In contrast to the absolute numbers of divergences as shown in Dietze et al. (2022) and Supplementary Figures S1-S3, the patients’ changes in lateralization patterns are not consistent within lesion groups. However, individual patients recovered especially with respect to one cluster of metrics, but showed poorer lateralization metrics in another cluster of Figure 5. One such example is patient S25, who had improved values in the metrics related to variability (cluster B) from the acute to the subacute phase, while showing poorer values for the metrics related to a shift of the auditory space (cluster A). The exact opposite behavior to S25 was shown by in S48. Obviously, only those metrics that were far from normal values in the first measurement phase can change by a large amount in the later measurements.

## 4 Discussion

The aim of this longitudinal study was to quantify the effects of ischemic stroke on binaural perception in a population of patients with only mild symptoms of stroke and with different lesion locations from the acute, to the subacute and the chronic phase of stroke. We hypothesized that binaural performance recovers toward later measurements.

In our population, binaural unmasking, assessed with a dichotic tone-in-noise detection experiment was not substantially affected by ischemic stroke and was constant over the measurement phases. For many patients, in contrast, the lateralization patterns differed from the control group in the acute phase of stroke. Towards the later measurements, many patients showed recovery of their lateralization abilities, which is in line with our hypothesis. This is comparable to other adaptive and maladaptive effects and to the spontaneous recovery observed after stroke as discussed in Cramer et al. (2011), and to the fast relearning of binaural hearing after alterations of binaural cues induced by a changed periphery (reviewed by Wright and Zhang, 2006). Recovery after stroke is linked to many factors, including experience. Butler (1987) showed that after altering the interaural cues, only those acoustic features that the participants had training in were relearned. Since this was not part of our investigations, we do not know how the patients behaved between the measurements, and the extent to which they trained everyday sound-source localization situations.

A smaller group of patients’ lateralization performance deteriorated over time. Deteriorated performance could be explained by secondary cell death of synaptically deprived brain areas, as suggested by Kolb and Teskey (2012). It could be also due to the imbalance of neuronal activity in the two hemispheres with hyperactivity in the contralesional hemisphere that could, in turn, further suppress the lesioned hemisphere by callosal inhibition and may reflect mal-plasticity, as suggested by Thompson et al. (2012).

### 4.1 Confounding Factors and Limitations of the Study

Studies in clinical populations pose simultaneous advantages and disadvantages (Gallun, 2021). A large number of participants can be measured when the experimental paradigm is adjusted to be as short as possible. Valuable information on the abilities of patients with clinically manifested impairments can be obtained, but is confounded by many factors. Even though we assessed the pure-tone hearing thresholds and cognitive state of the patients, these are possible confounding factors whose influence cannot be removed from the results. Hearing loss, and especially asymmetric hearing loss, poses challenges to interpreting the lateralization patterns. Hearing-threshold asymmetry was correlated with lateralization metrics associated with shifts of the lateralization pattern. However, it was not part of our measurements to assess the causes of the participants’ hearing loss. It is also not clear how the general state of the participants differed across the three measurements, but with the lateralization patterns being worst in the acute phase, their cognitive capacities were also lowest at this measurement. On the other hand, the motivation to participate in the study might have differed for the three measurement phases for each patient individually and was not assessed. The high dimensionality of this data set and the variety of lesion locations, hearing losses, and cognitive capacities, complicate the interpretation of the data. Yet even in the chronic phase, 6 of 22 patients still clearly showed impaired binaural abilities (at least 8 divergences from the control group), despite mild or absent clinically registered symptoms of stroke.

The acute phase measurements of this study were carried out on 50 patients, but only a subset of them participated again in the later measurement phases. Consequently, the number of longitudinally assessed patients was reduced to 31 who participated in more than one measurement. Although this number of participants is comparable even to other single-appointment studies on this topic (e.g., 21 in Bamiou et al., 2012; 22 in Aharonson et al., 1998; 50 in Spierer et al., 2009), it is a limitation. With more participants, the different lesion-location groups could be better represented, and confounding factors would be more evenly distributed across these groups.

### 4.2 Advantages of cue-specific experiments

As mentioned above, re-learning to use altered binaural information relies on exposure to the specific acoustic features (Butler, 1987). ILD or ITD cues in isolation are not present in real-life listening scenarios. Instead, each cue is accompanied by the matching other cue as well as by spectral information. In the frequency range of the stimuli used here (centered at 500 Hz), listeners usually rely on ITD-cues, whereas for far-field sources, ILDs are negligible at these low frequencies (Strutt, 1907). In the current study, one cue is always fixed at zero, while the other varies, resulting in artificial combinations of ITD and ILD. In some patients, we observed that the lateralization of ITD stimuli recovered better than the lateralization of ILD stimuli (e.g., S19, S25, S32, S36, S40). However, the opposite was also found in some patients (S19, S41). It is important to note that stimuli with ILD or ITD were presented in an interleaved fashion, ensuring that the same reference system was used.

We showed that headphone-based independent manipulation of interaural cues facilitates the detection of binaural-processing impairments that may remain undetected in localization (i.e. loudspeaker identification) experiments. This is most likely because real-world cues with high redundancy in the interaural cues are presented in localization tasks. After 40 years of research on cortical spatial maps, Middlebrooks (2021) concluded that unlike for other sensory modalities, no cortical spatial map exists for the auditory system. Instead, auditory space is represented by highly dynamic spatial neurons in the cortex (Middlebrooks, 2021). Consequently, it is possible that these neurons can react dynamically to altered binaural information, leading to a complete recovery of spatial hearing in free-field localization tasks, despite no recovery or even maladaptive effects for the artificial headphone-stimuli presented here. These stimuli are not externalized and not experienced in real-life listening scenarios, nor do they contain redundant information. Although more ecologically valid tasks can inform better about direct consequences in everyday life, cue-specific tasks as used in our study uncover difficulties in the underlying basic processing.

### 4.3 Relevance of binaural impairments to daily life

The benefits that arise from binaural hearing are undeniably important in everyday life (Avan et al., 2015), and binaural tests have been suggested to capture the variability across listeners with auditory difficulties that is not associated with classical monaural auditory tests such as pure-tone audiometry (Diedesch et al., 2021). Therefore, the influences of impaired binaural hearing are of relevance to anyone dealing with stroke. Affected individuals may or may not be aware of an existing impairment in binaural hearing. Patient-reported difficulties in spatial hearing after stroke were shown in Bamiou et al. (2012), whereas Javer and Schwarz (1995) reported that participants were not aware of their localization bias. Due to time limitations, we did not systematically investigate patient-perceived impairments.

We showed that some patients’ ability to use binaural information for the lateralization task recovered, whereas for others no recovery was observed until the last measurement. In both groups, we can assume that capacities are spent on either adaptive processes or managing the impairments in daily life, resulting in a higher listening effort in situations where binaural hearing is exploited. The additional cognitive load caused by stroke-induced impairments in binaural hearing “can interfere with other operations such as language processing and memory for what has been heard” (Peelle, 2017).

At the same time, studies such as ours, that determine the conditions under which recovery of perception occurs, “can provide insight into the plasticity and structure of the underlying neural processes. They can also inform the extent to which, and how best, individuals with impaired sound-localization abilities can be aided through training.” (Wright and Zhang, 2006). A study on the common stroke-induced phenomenon of neglect showed that deficits in spatial perception in different modalities were reduced by auditory spatial stimuli (Kaufmann et al., 2022). Rehabilitation training should therefore also rely on training in the auditory domain for relearning of auditory and non-auditory spatial perception, such as the positive effects of music listening on general recovery after stroke that were demonstrated by Särkämö et al. (2010). Carlile (2014) similarly pointed out the relevance of multi-modal training. He showed that compared to visual inputs, the involvement of the motor-state is even more important for the capacity to recalibrate to acoustic cues.

## 5 Conclusion

In this study, the effects of ischemic stroke on binaural perception were quantified using longitudinal measurements in the acute, subacute, and chronic phases of stroke in a population of patients having only mild symptoms and with different lesion locations. We found that binaural hearing abilities are impaired in many patients, and that the severity of the impairment changes over time after stroke onset. While many patients’ lateralization abilities recovered toward later measurements, deteriorated performance was observed for others. Since stroke is such a common medical condition, its effect on binaural hearing should be investigated more thoroughly. The insights gained during this study can guide future research with respect to the management of confounding factors and to the relevance of choosing experimental conditions that best uncover impaired processing. Identifying which medical conditions lead to impaired binaural hearing might not only help in designing effective rehabilitation programs, but should also be communicated to the patients.

## 7 Author Contributions

AD, MD, and PS contributed to the conception and design of the study. AD, MD, PS, and KW planned the experimental procedures. AD organized the database and performed analysis. AD and MD interpreted the data. AD wrote the first draft of the manuscript. MD and HP wrote further sections of the manuscript. All authors contributed to manuscript revision, read, and approved the submitted version.

## 8 Acknowledgements

We thank Matthias Bröer and Anna Methner for their help with data collection and Prof. Dr. Claas Unverferth for his support in scheduling appointments and providing a room for the subacute phase measurements.

## Supplementary Material

**Figure S 1.**
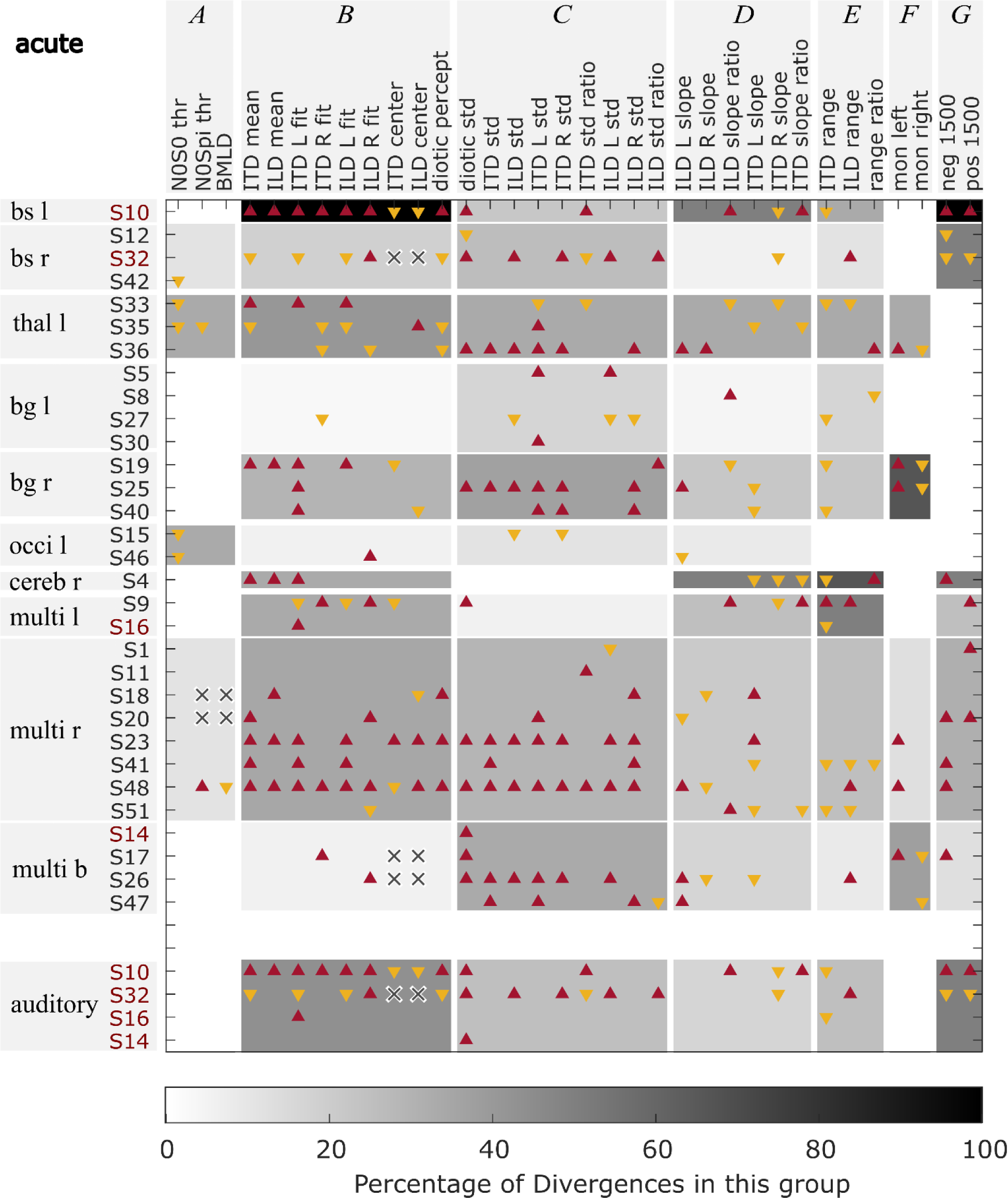
Divergences in the acute phase measurements from the control group for different variables. Red upward triangles indicate values of individuals that are larger and yellow downward triangles values that are smaller than the normal values (mean ±1.5 times standard deviation) calculated from the control group. Crosses indicate missing values. The gray shading indicates the percentage of deviations found within one lesion group for one of the variable groups (A–G). The red font is used for those patients who had a lesion on the primary auditory pathway.

**Figure S 2.**
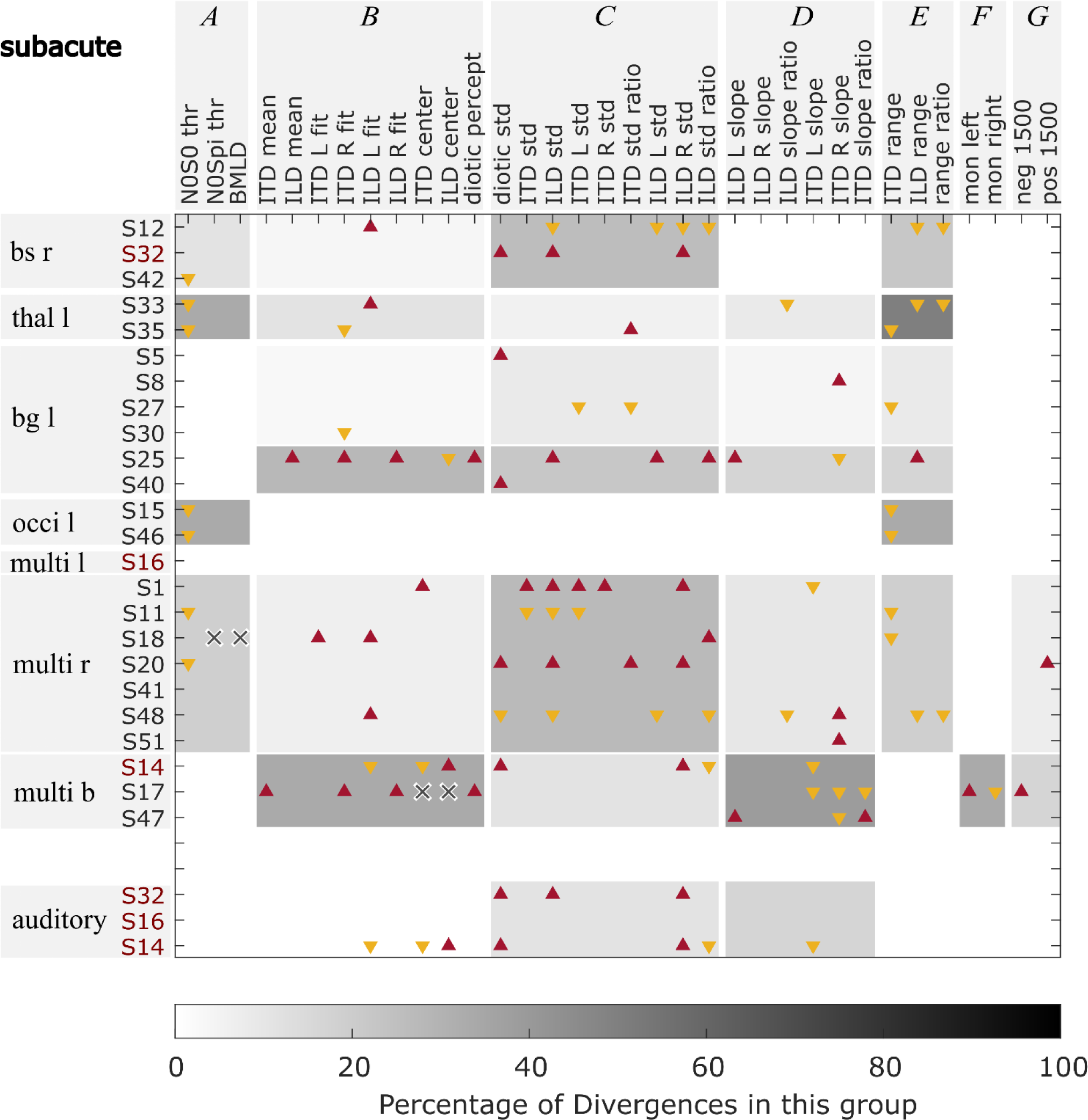
Divergences in the subacute phase measurements from the control group for different variables. Red upward triangles indicate values of individuals that are larger and yellow downward triangles values that are smaller than the normal values (mean ±1.5 times standard deviation) calculated from the control group. Crosses indicate missing values. The gray shading indicates the percentage of deviations found within one lesion group for one of the variable groups (A–G). The red font is used for those patients who had a lesion on the primary auditory pathway.

**Figure S 3.**
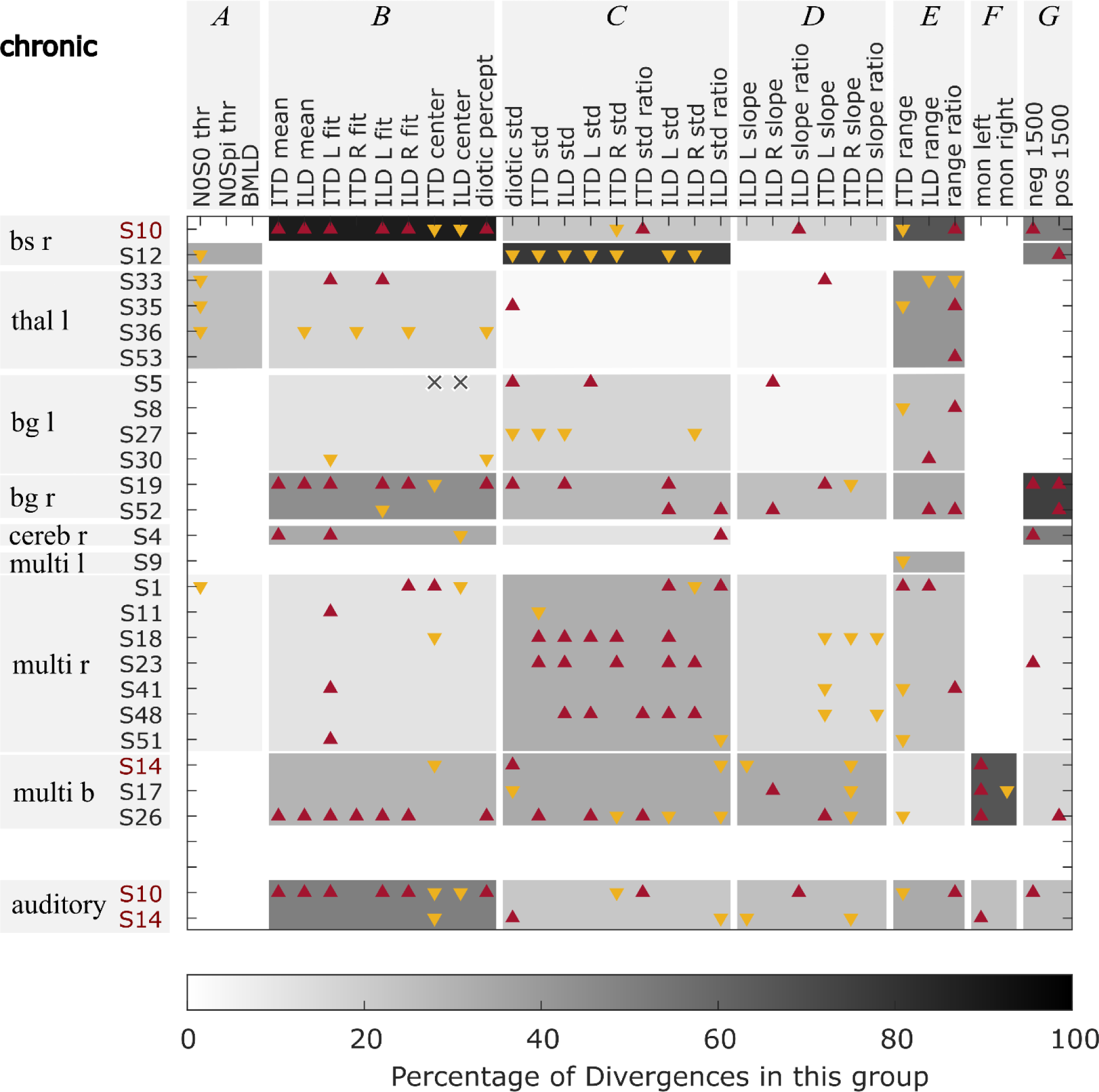
Divergences in the chronic phase measurements from the control group for different variables. Red upward triangles indicate values of individuals that are larger and yellow downward triangles values that are smaller than the normal values (mean ±1.5 times standard deviation) calculated from the control group. Crosses indicate missing values. The gray shading indicates the percentage of deviations found within one lesion group for one of the variable groups (A–G). The red font is used for those patients who had a lesion on the primary auditory pathway.

